# Investigation of cellular and molecular changes linked with neuropathic pain in healthy and injured human trigeminal nerves

**DOI:** 10.1101/2024.10.05.616798

**Authors:** Martina Morchio, Ishwarya Sankaranarayanan, Diana Tavares-Ferreira, Natalie Wong, Simon Atkins, Emanuele Sher, Theodore J Price, Daniel W Lambert, Fiona M Boissonade

**Affiliations:** Neuroscience Institute and School of Clinical Dentistry, University of Sheffield, Sheffield, UK; Department of Neuroscience and Center for Advanced Pain Studies, University of Texas at Dallas, Richardson, TX, USA; Eli Lilly and Company, Lilly UK Neuroscience Hub, Bracknell, UK

## Abstract

Injuries to the trigeminal nerve, responsible for sensory innervation to the face, may occur during routine dental procedures, resulting in the formation of a neuroma accompanied by loss of sensation and/or symptoms of pain. In order to gain insight into the molecular mechanisms underpinning the sensory changes, single nuclei RNA sequencing and spatial transcriptomics were employed to profile the transcriptional landscape at single cell resolution of human trigeminal nerves and neuromas. Cellular and transcriptional changes were identified that correlated with the presence of pain, including an expansion of endothelial cells with a pro-inflammatory phenotype and over-expression of *HLA-A*, *CXCL2* and *CXCL8*. Interactome analysis highlighted signalling changes linked with the presence of pain. HLA-A protein expression was confirmed in neuromas and positively correlated with symptoms of pain. The atlas generated represents a valuable resource for pain research, highlighting the role of inflammation, endothelial cell dysfunction and chemokine signalling in neuropathic pain.

## Introduction

Approximately 300,000 peripheral nerve injuries arise each year in Europe^1^. In addition to loss of motor and sensory function, a proportion of patients have persistent pain for which there is no reliable treatment^1^. This is associated with sleep disturbances, depression and various debilitating psychological problems^2^. Injuries to branches of the trigeminal nerve, most commonly the lingual and inferior alveolar nerves, can occur as a result of routine dental procedures^3^. While peripheral nerves regenerate spontaneously, in some patients the presence of a gap between the proximal and distal ends, bone fragments, scar tissue and inflammation may prevent functional reinnervation of the target areas, resulting in the formation of a swollen mass termed neuroma^4^. The frequency of lingual nerve injuries during oral and maxillofacial procedures varies between 0.6-2%, resulting in anaesthesia, paraesthesia or hyperesthesia of the floor of the mouth, the lingual gingiva and the anterior part of the tongue, affecting everyday activities such as speaking, eating and drinking and potentially leading to altered taste^5–7^. In addition, some patients report the presence of neuropathic pain, often described as a burning sensation^6^. To treat these symptoms, patients can undergo nerve repair surgery where the neuroma is resected and the nerve ends are surgically reconnected, promoting functional recovery of sensation^6^.

Human lingual neuromas represent a unique resource to investigate mechanisms linked with neuropathic pain. The comparison between samples from patients that have incurred the same type of injury, where some, but not all, report symptoms of pain, enables the identification of factors specifically linked with neuropathic pain, which are independent from the pathophysiological changes associated with nerve injury and regeneration. Previous work on human neuromas highlighted changes in the molecular expression of selected targets including ion channels, MAP kinases and inflammatory mediators^8,9^. High-throughput bulk transcriptomics has been employed to identify overall differences in non-trigeminal neuromas compared to healthy uninjured controls^10^. However, detailed characterisation of the cellular composition and the transcriptional changes linked with the severity of pain at single cell level in human neuromas has never been performed.

In this work, we sought to characterise the cellular and transcriptional composition of healthy and injured trigeminal nerves and identify molecular changes in human trigeminal neuromas linked with the presence of pain. Single nucleus RNA sequencing was employed on samples of healthy and injured human trigeminal nerves to characterise the cellular composition at single cell resolution. Spatial transcriptomics was performed on a larger pool of human lingual neuromas, including both painful and non-painful samples, to characterise the transcriptional landscape within the morphological context and identify changes in gene expression localised within the nerve fascicles. Cell-cell interactome analysis was performed to identify changes in signalling networks linked with the presence of pain. RNAscope and immunohistochemistry were used to validate the findings.

Several transcriptional changes linked with the presence of pain were identified, including over-expression of *HLA-A, CXCL2* and *CXCL8* in painful samples, highlighting the role of inflammation in neuropathic pain. The protein expression of *HLA-A* was confirmed in neuromas and positively correlated with symptoms of pain. The unique atlas generated here provides a detailed spatial overview of the cell types that populate human peripheral trigeminal nerves, in health and injury, and their transcriptional profile. The comparison of painful and non-painful samples highlighted several changes in cellular composition, transcriptome and inferred molecular signalling linked specifically with the presence of pain.

## Results

### Single nuclei RNA sequencing reveals the cellular composition of healthy and injured human trigeminal nerves

In order to characterise the cellular composition of trigeminal nerves with and without injury in humans, single nuclei RNA sequencing was employed on samples of mechanically dissociated trigeminal nerves and neuromas. Trigeminal nerves were obtained from the Netherland Brain Bank from two donors who died of non-neurological causes and without a diagnosis of dementia (Table 1). Lingual nerve neuromas were obtained from lingual nerve repair surgeries carried out at the Royal Hallamshire Hospital in Sheffield, UK (Table 2).

**Table 1.**
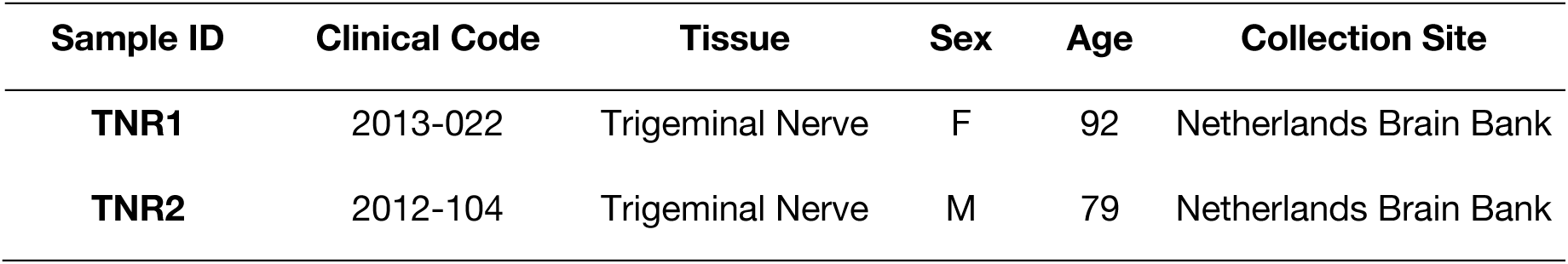
Information linked to the samples of trigeminal nerve roots used for single nuclei RNA sequencing. The trigeminal nerve root samples were obtained from the Netherland Brain Bank from healthy organ donors without diagnosis of other neurological conditions.

| **Sample ID** | **Clinical Code** | **Tissue** | **Sex** | **Age** | **Collection Site** |
| --- | --- | --- | --- | --- | --- |
| **TNR1** | 2013-022 | Trigeminal Nerve | F | 92 | Netherlands Brain Bank |
| **TNR2** | 2012-104 | Trigeminal Nerve | M | 79 | Netherlands Brain Bank |

**Table 2.** Clinical information and applications of the human neuroma samples. The lingual neuromas were obtained during nerve repair surgeries performed at the Royal Hallamshire Hospital in Sheffield, UK.

| Sample ID | Time since injury (months) | Age | Sex | Pain VAS | Spatial | snRNAseq | IHC |
| --- | --- | --- | --- | --- | --- | --- | --- |
| LN1 | 12 | 27 | F | 55 | Yes |  | Yes |
| LN2 | 7 | 41 | M | 2 | Yes |  |  |
| LN3 | 20 | 29 | F | 0 |  |  | Yes |
| LN4 | 19 | 65 | F | 100 |  |  | Yes |
| LN6 | 14 | 25 | F | 0 |  |  | Yes |
| LN7 | 17 | 30 | F | 4 | Yes |  | Yes |
| LN8 | 4 | 36 | M | 14 | Yes |  | Yes |
| LN10 | 69 | 39 | F | 0 |  |  | Yes |
| LN11 | 18 | 33 | F | 3 |  |  | Yes |
| LN12 | 16 | 34 | F | 43 | Yes |  | Yes |
| LN13 | 4 | 28 | F | 65 | Yes | N2 | Yes |
| LN14 | 13 | 31 | F | 0 |  | N1 |  |
| LN15 | 12 | 34 | F | 61 | Yes |  | Yes |
| LN18 | 17 | 46 | F | 72 |  |  | Yes |

A total of 86,831 nuclei were sequenced: 54,144 from two trigeminal nerve samples and 32,687 from two neuroma samples. Ambient RNA was removed with CellBender and doublets were removed using scDblFinder, removing nuclei with a doublet score higher than 0.5. Nuclei with fewer than 500 reads and 250 genes detected, a novelty score lower than 0.8 and the presence of mitochondrial genes higher than 5% were removed. After quality control, data from 56,959 nuclei were kept, 21,990 from lingual neuromas and 34,969 from trigeminal root nerves, with a median of 1,305 genes detected per nucleus. The data was analysed with Seurat, performing SCT normalisation, rPCA integration and clustering at resolution 0.5. A total of 26 clusters were identified with annotation markers derived from the literature (Supplementary table 1)^11–15^ (Figure 1.B,C). In the following paragraphs, the clusters identified by snRNA-seq are described in more detail.

**Figure 1.**
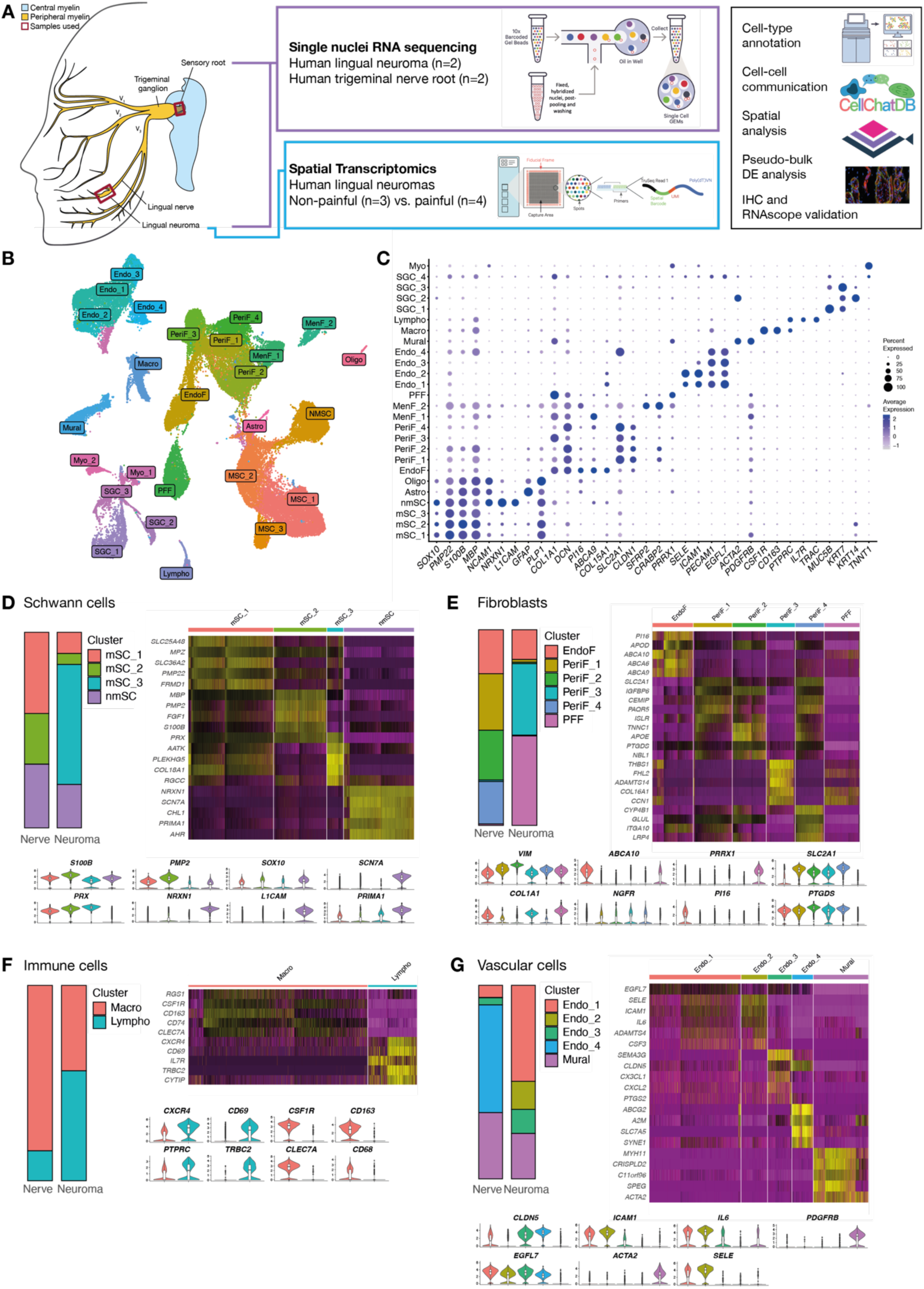
Overview of single nuclei RNA sequencing atlas. A: Summary of the samples and methods used to characterise the cellular and transcriptional composition of healthy and injured lingual nerves and identify factors associated with neuropathic pain. B: UMAP of nuclei from human trigeminal nerve roots and lingual neuromas analysed by single nuclei RNA sequencing. C: Dotplot displaying marker genes used to annotate the cell types in human trigeminal nerves. D-G: Overview of the major cell-types populating trigeminal nerves and neuromas: Schwann cells (D), fibroblasts (E), vascular (F) and immune (G). For each major cell type, a barplot displays the proportion of each cell subtype in trigeminal nerves and neuromas, a heatmap displays the top marker genes, where yellow indicates a higher level of expression, and violin plots displaying the expression levels of other genes used for annotation.

### Glial cells in human trigeminal nerves and neuromas

Three clusters were annotated as myelinating Schwann cells: mSC_1, mSC_2 and mSC_3. mSC_1 and mSC_2 are found in both neuromas and trigeminal nerve root samples and are enriched in markers for myelination such as *PMP22, MBP* and *MPZ* (Figure1.D). A third myelinating Schwann cell cluster, mSC_3, was found only in the neuroma samples. mSC_3 is enriched in markers for myelination (*PRX*) but also genes indicating stress (*AATK*, involved in apoptosis^16^), inflammation (*TNFRSF25*, member of the TNF family that induces the activation of NF-kB pathway^17^) and keloid formation (*COL18A1*, involved in ECM deposition and more highly expressed in keloidal Schwann cells compared to healthy skin Schwann cells^18^). Non-myelinating Schwann cells (nmSC) were detected in both neuromas and trigeminal nerve roots and display enrichment in genes previously identified as nmSC markers including *NRXN1, SCN7A* and *PRIMA1*^19^.

Yim, et al. ^19^ identified a myelinating Schwann cell subpopulation in mouse and human nerves that expresses high levels of *PMP2* and preferentially myelinates large diameter axons, in particular motor axons, which might correspond to the mSC_2 cluster identified here, where *PMP2* expression is significantly upregulated. While motor axons would be found in the motor root of the trigeminal nerve, other large-diameter axons such as Aβ low threshold mechanoreceptors present in the sensory root of the trigeminal nerve might be preferentially myelinated by the mSC_2 cluster.

Central glial cells were identified in the trigeminal nerve roots (Supplementary Fig.A), where one cluster characterised by *GFAP* expression was annotated as astrocytes (Astro) and one positive with *MOG* and *OLIG1* as oligodendrocytes (Oligo). These clusters were absent in the neuromas and are specifically abundant in one trigeminal nerve root sample, possibly due to the inclusion of the peripheral to central transition zone where peripheral myelin is replaced by central myelin^20^.

### Fibroblast heterogeneity in human trigeminal nerves

Among the fibroblast subtypes, endoneurial (EndoF), perineurial (PeriF_1-4) and profibrotic (PFF) fibroblasts were detected in both sample types (Figure 1.E), while in the trigeminal nerve roots, meningeal fibroblasts were also identified (MenF_1-2) (Supplementary Fig 1.A). All clusters robustly express the general fibroblast marker vimentin (*VIM*) (Figure 1.E). The endoneurial cluster, found in both neuroma and healthy nerve samples, was enriched in *PI16*, *ABCA6, 9* and *10* and *APOD*. PI16 was identified by Singhmar, et al. ^21^ to promote pain-like behaviour following sciatic nerve injury in rats by inducing permeability of the blood-nerve barrier and increasing immune cell infiltration; however, in the rat sciatic nerve PI16 expression is localised to the perineurial and epineurial layer^21^. In contrast, immunohistochemistry confirmed PI16 localization to endoneurial-like cells in human neuromas (Figure 2.A).

**Figure 2.**
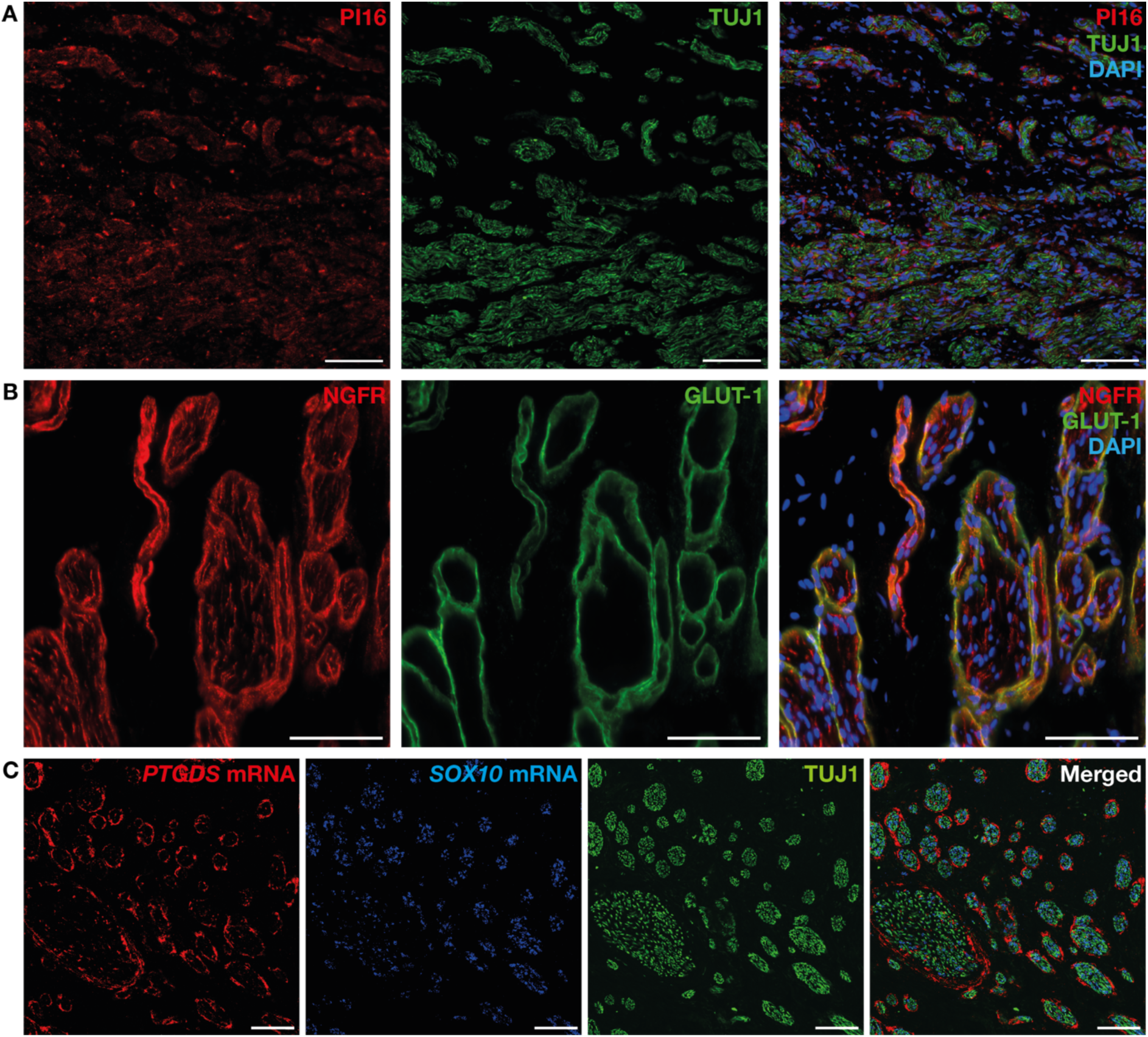
Protein and RNA expression of fibroblast marker genes in human neuromas. A: Representative image of PI16 immunolabelling in endoneurial-like cells within nerve fascicles, labelled by TUJ1. B: Representative image of dual-labelling of the p75 NGF receptor (NGFR) and GLUT-1, a marker for perineurial cells, showing co-localization. C: Representative image of *PTGDS* and *SOX10* RNAscope combined with TUJ1 immunolabelling, highlighting the location of *PTGDS* mRNA in perineurial-like structures surrounding nerve fascicles. All scale bars are 100 µm. Images taken at 40X magnification.

Perineurial fibroblast clusters are found in varying proportions across sample types: PeriF_1,2 and 4 were largely found in the trigeminal root samples, while PeriF_3 was found primarily in the neuroma samples. PeriF_1,2,4 were enriched in genes such as *IGFBP6* and *SLC2A1*, widely used as a perineurial marker gene^22^. PeriF_3 displayed increased expression of *THBS1, CCN1* and *ADAMTS14.* THBS1 is secreted by fibroblasts in peripheral nerves following transection injury, promoting neurite outgrowth^23^. CCN1 promotes SC proliferation and upregulation of c-Jun^24^. *NGFR* was expressed in the perineurial clusters to a higher level compared to Schwann cells, validated at the protein level with immunohistochemistry (Figure 2.B), confirming previous reports in rat perineurial cells^25,26^. *PTGDS*, encoding for prostaglandin D2 synthase, was also highly expressed in perineurial fibroblasts, confirmed by RNAscope analysis, highlighting its location to perineurial structures surrounding TUJ1+ fibres and *SOX10*+ Schwann cells (Figure 2.C). PTGDS is an enzyme involved in the conversion of PGH_2_ to PGD_2_, leading to neuronal sensitization and pro-nociceptive effects in both animal models and humans^27–29^.

Finally, profibrotic fibroblasts (PFF) were particularly abundant in the neuromas, displaying high expression of the profibrotic marker *PRRX1*^30^. This cluster was enriched in ECM-related genes including *COL1A1, FBLN1, COL3A1* and *COL1A2,* as well as inflammation-related genes such as *FOSB,* involved in promoting a profibrotic programme in pulmonary fibrosis^31^ and *LSP1,* which mediates neutrophil activation^32^.

### Vascular cell expansion in human neuromas

Endothelial cells, vascular smooth muscle cells and pericytes make up the vasculature that supplies peripheral nerves. A total of 9,113 nuclei were annotated as vascular, including four clusters of endothelial cells (Endo_1, Endo_2, Endo_3, Endo_4) and one cluster of mural cells, containing vascular smooth muscle cells and pericytes (Figure 1.G). In the neuromas, a higher proportion of endothelial cells were identified, making up 30% of the nuclei in those samples, while only 3% of the nuclei in the nerves were annotated as endothelial, the majority of which belong to the Endo_4 cluster.

All the endothelial cell clusters expressed high levels of canonical endothelial cell markers such as *EGFL7*^13^. Additionally, Endo_1 was enriched with genes usually found in cytokine activated endothelial cells and involved in leukocyte recruitment at the site of injury, such as *SELE*^33^, *ICAM-1*, considered a master regulator of inflammation and injury resolution^34^ and interleukin-6 (*IL6*), a hallmark of inflammation arising following nerve injury^35^. Endo_4, only found in the trigeminal nerve roots, displayed high expression of genes involved in blood-brain barrier formation such as *SLC7A5*^36^.

The mural cell cluster expressed *ACTA2, MYH11* and *PDGFRB*, markers for both vascular smooth muscle cells and pericytes^37^. In keeping with a report by Muhl, et al. ^37^, mural cells displayed less heterogeneity than fibroblasts, despite the marked morphological differences between pericytes and vascular smooth muscle cells.

### Immune cell populations in trigeminal nerves and neuromas

Two immune cell subpopulations were identified: lymphocytes (602 nuclei) and macrophages (2,204 nuclei) (Figure 1.F). The macrophage cluster expressed high levels of *RGS1*, marker for both macrophages and T-cells^38,39^, *CSF1R* and *CLEC7A*, commonly used as specific macrophage markers^14,40^ and *CD163*, a marker for M2 macrophages^41^.

The lymphocyte cluster displayed high levels of *CXCR4* and *CD69*, T-cell markers^42^, *CYTIP*, also found to be expressed in T cells^43^ and *TRBC2*, involved in antigen binding activity^44^. In the neuroma samples a higher proportion of lymphocytes are detected compared to macrophages. T-cells have been identified to have an important role in nerve repair, modulating remyelination and inflammation^45^.

Finally, four subtypes of salivary gland cells (SGC_1, SGC_2, SGC_3, SGC_4) and one cluster of myocytes (Myo) were identified specifically in the neuroma samples (Supplementary Fig 1.B)

### Spatial transcriptomics identifies distribution of cell types within the morphological context of human neuromas

Visium spatial transcriptomics was employed to characterise the spatial distribution of cell types and the transcriptional signature in the morphological context of the tissue. Consecutive sections from 7 human neuroma samples (Table 2) were placed on Visium slides and processed for spatial transcriptomics. The data was analysed with SpaceRanger and Giotto, using Harmony to integrate the data and Leiden clustering to identify subpopulations. A total of 17 clusters were identified and annotated based on the genes identified from snRNAseq and the literature^11–15^. Spots enriched for fibroblasts, endothelial cells, Schwann cells, perineurial cells, myocytes, B cells and macrophages were annotated based on the top differentially expressed genes (Supplementary table 2). The heatmap in Figure 3.A displays the top marker genes for each cluster, while representative sections from a painful sample (LN15) and a non-painful one (LN2) are shown in Figure 3.B,C.

**Figure 3.**
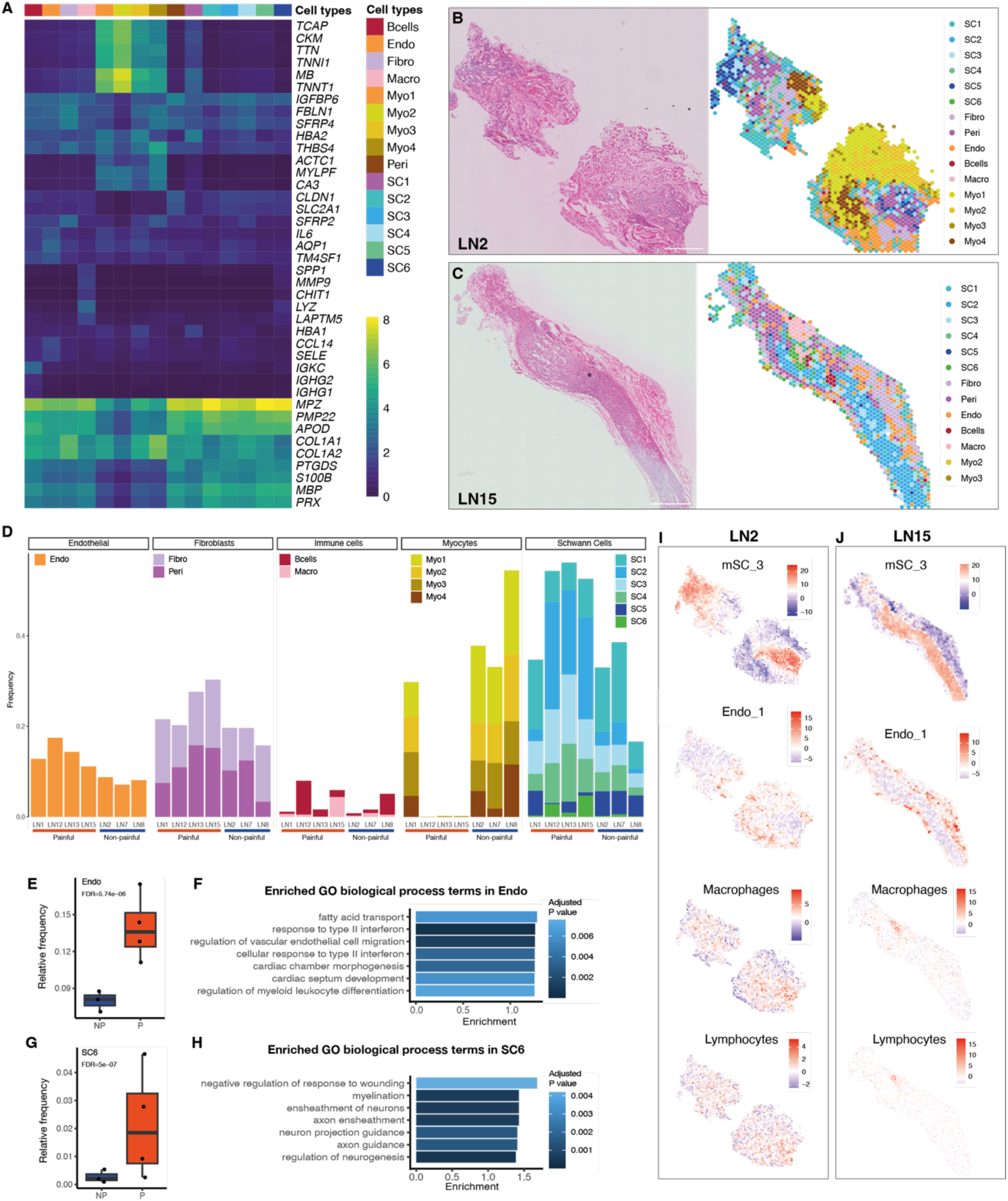
Spatial transcriptomics of human neuromas. A: Heatmap displaying the expression of marker genes for each cell type identified by Visium analysis in human lingual neuromas where yellow indicates higher expression. B-C: Representative sections of LN2 (pain VAS: 2) and LN15 (pain VAS: 61) stained with H&E (right) and corresponding spots colour-coded by the enriched cell-type identified by clustering. D: Enriched cell-type proportion across samples, divided by major cell type: from left to right, endothelial cells (Endo), fibroblasts (Fibro and Peri), immune cells (Bcells and Macro), myocytes (Myo1-4) and Schwann cells (SC1-6). E,G: Boxplot displaying the frequency of endothelial cells (E) and one subtype of Schwann cells (SC6, G) in painful and non-painful samples, where a statistical significant difference was identified with EdgeR and corrected by false discovery rate. F,H: Enriched GO terms for biological processes in differentially expressed genes in the Endo (F) and SC6 (H) cluster, where the x-axis indicates the enrichment, calculated by dividing the number of significant genes over the expected number of genes in each GO term. I,J: Representative images of PAGE analysis, where marker genes of the clusters annotated from the snRNA-seq analysis of human neuromas were used to identify the enrichment of specific cell types in the spatial data. Relative enrichment of cell-type gene signatures is displayed on a scale from lowest (purple) to highest (red), where each image has a separate scale bar. The enrichment of gene signatures characteristic of myelinating Schwann cells (mSC_3), endothelial cells (Endo_1), macrophages and lymphocytes is shown for LN2 (I) and LN15 (J).

The clusters were differentially distributed across samples (Figure 3.D). Schwann cell clusters (SC1-6), characterised by the expression of myelination genes such as *PMP22, MBP, PRX* and *S100B*, were the most abundant in LN12, LN13 and LN15. SC1 displays the expression of both Schwann cell marker genes and haemoglobin genes (*HBA1, HBA2*). Due to its localisation to the outer edge of tissue sections, the expression of haemoglobin transcripts might indicate the presence of residual red blood cells following surgical removal of the sample. The other five clusters were found across all samples mostly overlaying areas containing nerve fascicles.

The most ubiquitous cluster expressed genes enriched in fibroblasts including *COL1A1, FBLN1* and *COL1A2*^37^. The cluster was localised to areas surrounding the nerve fascicles, suggesting the presence of fibrotic tissue. This cluster displayed over-expression of *SFRP2* and *4*, encoding for secreted frizzled-related proteins that act as extracellular regulators of the Wnt signalling pathway by competing for Wnt ligand binding^46^. The Wnt pathway has been identified as a critical mediator of fibrosis, contributing to fibroblast activation and differentiation into myofibroblasts^47^. Changes in Wnt gene expression have been reported in both human and animal models of peripheral nerve injury, including *SFRP4* over-expression^48^.

The second most ubiquitous cluster (Endo) expressed *AQP1, TM4SF1* and *SELE*, indicating the presence of endothelial cells^13,49,50^. *IL6* and *CCL14* were also enriched in this cluster suggesting a pro-inflammatory phenotype of the endothelial cells present in these samples: the role of IL-6 as a pro-inflammatory molecule is well-known ^51^, while CCL14 has been shown to bind to CCR1, 3 and 5, potentially inducing leukocyte infiltration^52^.

The perineurial cluster (Peri) was enriched in *PTGDS, CLDN1* and *SLC2A1*^13,22^, and found across all samples in areas associated with nerve fascicles. In samples LN15 and LN12, which display a more ordered fascicular organization compared to the other samples, this cluster is distributed along the edges of the nerve fascicles, in line with the expected morphology of the perineurial barrier.

Four clusters (Myo1-4) enriched in myocyte markers, such as *MB, TNNT1, CKM* and *ACTC1*, were particularly abundant in LN2 and LN8. These clusters overlay areas with a morphology typical of skeletal muscles, characterized by the presence of multinucleated myocytes, and are not directly associated with nerve fascicles.

Finally, two clusters were enriched in immune cell markers. The cluster annotated as B cells (Bcells) was enriched in *IGKC*, encoding the Immunoglobulin Kappa Constant and *IGHG1-2*, encoding the immunoglobulin heavy constant gamma 1 and 2, respectively^53^. This cluster was found across all samples but was particularly abundant in LN12 and LN8. The cluster annotated as Macro was enriched in genes such as *LYZ*, a marker for macrophage activation^54^, *SPP1*, encoding a protein secreted by activated macrophages^55^, *LAPTM5*, associated with pro-inflammatory activation of macrophages^56^ and *CHIT1,* involved in macrophage polarization and activation^57^. The macrophage cluster also expressed high levels of *MMP9*, known to be secreted by Schwann cells in the early stages following nerve injury, promoting blood-nerve barrier breakdown and leukocyte influx at the site of injury^58^. The macrophage cluster was particularly abundant in LN15 (Figure 3.C), one of the samples classified as painful based on the patient’s reported VAS score (Table 2).

The location of specific cell types was confirmed using cell-type deconvolution with PAGE^59^. Examples are shown in Figure 3.I-H where the enrichment of mSC_3, Endo_1, the macrophage and lymphocytes gene expression signatures, obtained from the snRNA-seq dataset, is shown across the barcodes of a representative section of LN2 (Figure 3.I) and LN15 (Figure 3.J).

Differential abundance of cell types between painful and non-painful samples was tested with edgeR^60,61^ (Supplementary table 3). Among the top differentially abundant clusters were SC6 (Figure 3.E) and Endo (Figure 3.G), both more abundant in the painful samples. The endothelial cell cluster was enriched for gene ontology (GO) biological processes including response to interferon II and regulation of vascular endothelial cell migration and myeloid leukocyte differentiation (Figure 3.F). The Schwann cell cluster SC6 was enriched for GO terms including negative regulation of response to wounding, axon guidance and regulation of neurogenesis (Figure 3.H).

### Pseudo-bulk differential expression analysis identifies changes in painful neuromas involved in chemokine signalling and antigen presentation

Pseudo-bulk differential expression analysis was performed by selecting the barcodes overlaying nerve fascicles based on H&E staining, in order to focus on pathologically-relevant areas that are more likely to contribute to the development and maintenance of neuropathic pain (Figure 4.A). Principal component analysis (Figure 4.B) displayed the separation between painful and non-painful samples. The analysis identified 59 differentially expressed (DE) genes (padj < 0.05) (Figure 4.C). Among the top DE genes (padj < 0.01) are *HLA-A*, an MHC Class I gene involved in antigen presentation, chemokines *CXCL2* and *CXCL8*, the gene *NID2* encoding for a basement membrane protein, *SLC52A3*, encoding for a riboflavin transporter, the metalloprotease-encoding gene *MMP19* and *ADGRG1*, encoding for a G-protein coupled receptor (Figure 4.C).

**Figure 4.**
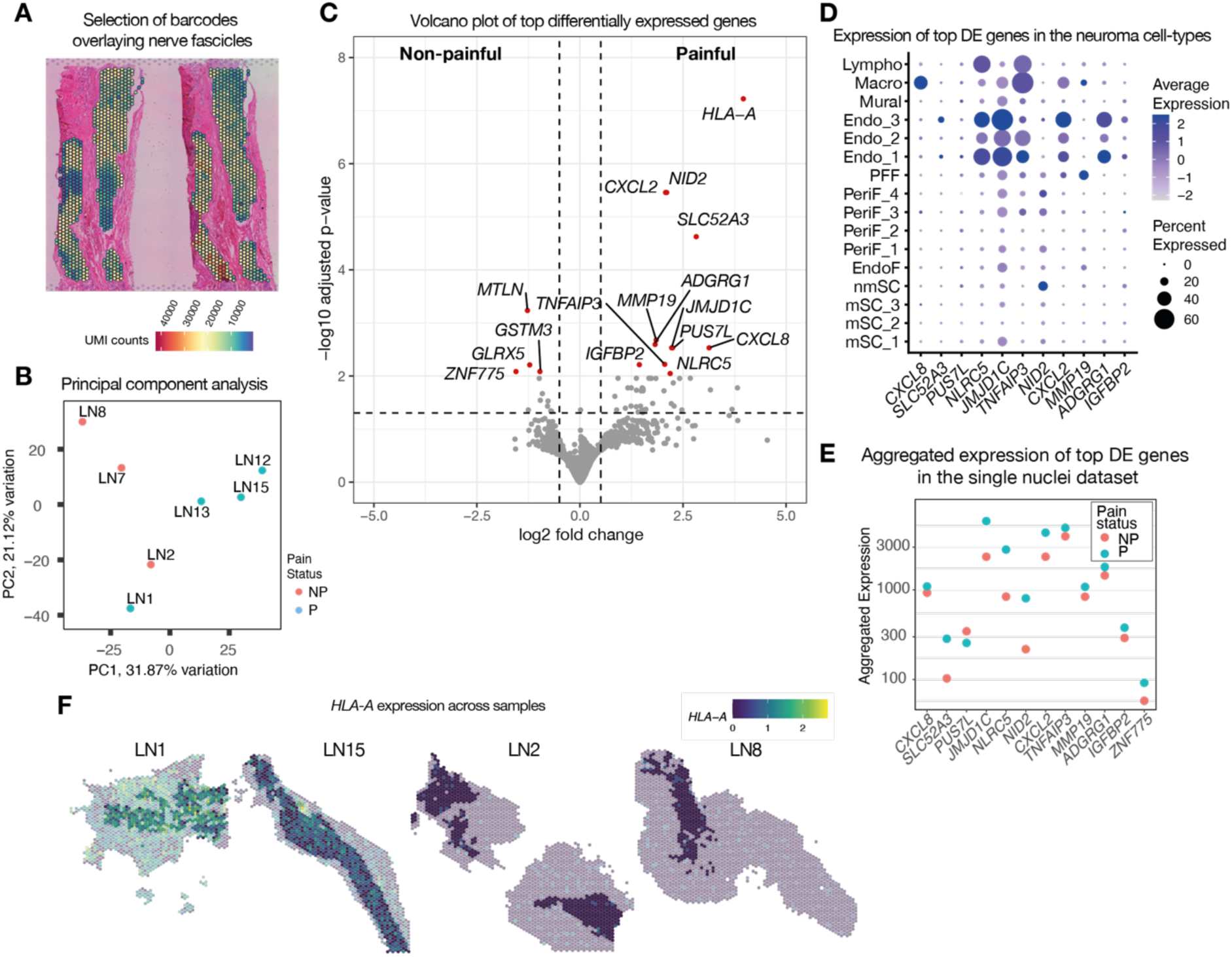
Pseudo-bulk differential expression analysis in painful and non-painful neuromas. A: Representative image displaying the selection of barcodes overlaying nerve fascicles in a sample of human neuroma. B: Principal component analysis (PCA) displaying the distribution of the samples used for pseudo-bulk DE analysis according to the first two principal components displayed on the x and y axes. C: Volcano plot displaying the top DE genes in painful and non-painful neuromas, where the x-axis represents the logarithmic fold change in expression and the y-axis represents the adjusted p-value. D: Dotplot displaying the expression of the top DE genes in the cell-types identified by snRNAseq of human neuromas. E: Plot representing the aggregated level of expression of top DE genes in the painful and non-painful samples analysed by snRNAseq in human neuromas. F: Representative images of the expression of *HLA-A* across four samples of human neuromas classified as painful (LN1 and LN15) and non-painful (LN2 and LN8), where the barcodes overlaying nerve fascicles used for DE analysis are shown in full colour, while the ones not in the fascicles are opaque, and all barcodes are colour-coded to indicate the expression level from low (blue) to high (yellow).

The expression of the top DE genes was confirmed in the single nuclei dataset for the neuromas (Figure 4.D), with the exception of *HLA-A*, as the probes for this gene were absent from the probe set used for snRNA-seq. Several of the DE genes were expressed in endothelial cells, particularly *NLRC5*, a trans-activator of *HLA-A*, *JMJD1C*, a histone demethylase, the chemokine *CXCL2, SLC25A3, ADGRG1* and *TNFAIP3*, encoding for a Zinc finger protein and involved in TNF-mediated apoptosis, also expressed in macrophages and lymphocytes*. CXCL8* was expressed in macrophages while *NID2* was expressed in non-myelinating Schwann cells. By comparing the expression of the DE genes in the samples used for single nuclei RNA sequencing analysis, which included one painful and one non-painful neuroma, the direction of the change in expression was confirmed in all genes except for *PUS7L*, a pseudourinidase synthase, which was upregulated in the painful samples as measured by spatial transcriptomics but displayed lower expression in the painful sample compared to the non-painful one in the snRNAseq dataset (Figure 4.E). The expression profile of *HLA-A* in representative sections of two painful and two non-painful samples is shown in Figure 4.F.

### Changes in cell-cell interactions in painful neuromas

In order to begin to understand the cell-cell interactions occurring in painful and non-painful neuromas, CellChat v2 was used on the combined spatial datasets grouped by pain status. CellChat v2 takes into account both the level of expression and spatial location of each ligand-receptor pair from a manually curated database^62^. The spots were subset and merged to include Schwann cells (SC), endothelial cells (Endo), fibroblasts (Fibro), perineurial cells (Peri), B cells (Bcells) and macrophages (Macro). In total, 8,350 interactions were inferred in painful samples while 7,509 interactions were inferred in non-painful ones (Figure 5.B). The postulated differences in information flow between painful and non-painful samples in pathways with significant interactions is shown in Figure 5.A. The change in cell-type contribution of signalling is shown in the chord plot in Figure 5.C-D, where line thickness indicates the number of interactions. Overall, the painful samples display increased connectivity among the macrophages, perineurial cells, Schwann cells and endothelial cells. Analysis of differentially expressed ligands and receptors, identified several ligand-receptor interactions upregulated in the painful samples, particularly involving endothelial cells (Figure 5.E), which display an increased likelihood to interact with perineurial cells through CLDN1, DHH and FGF1 signalling, with Schwann cells through MHC class II molecules and with endothelial cells through the activation of the LIF receptor.

**Figure 5.**
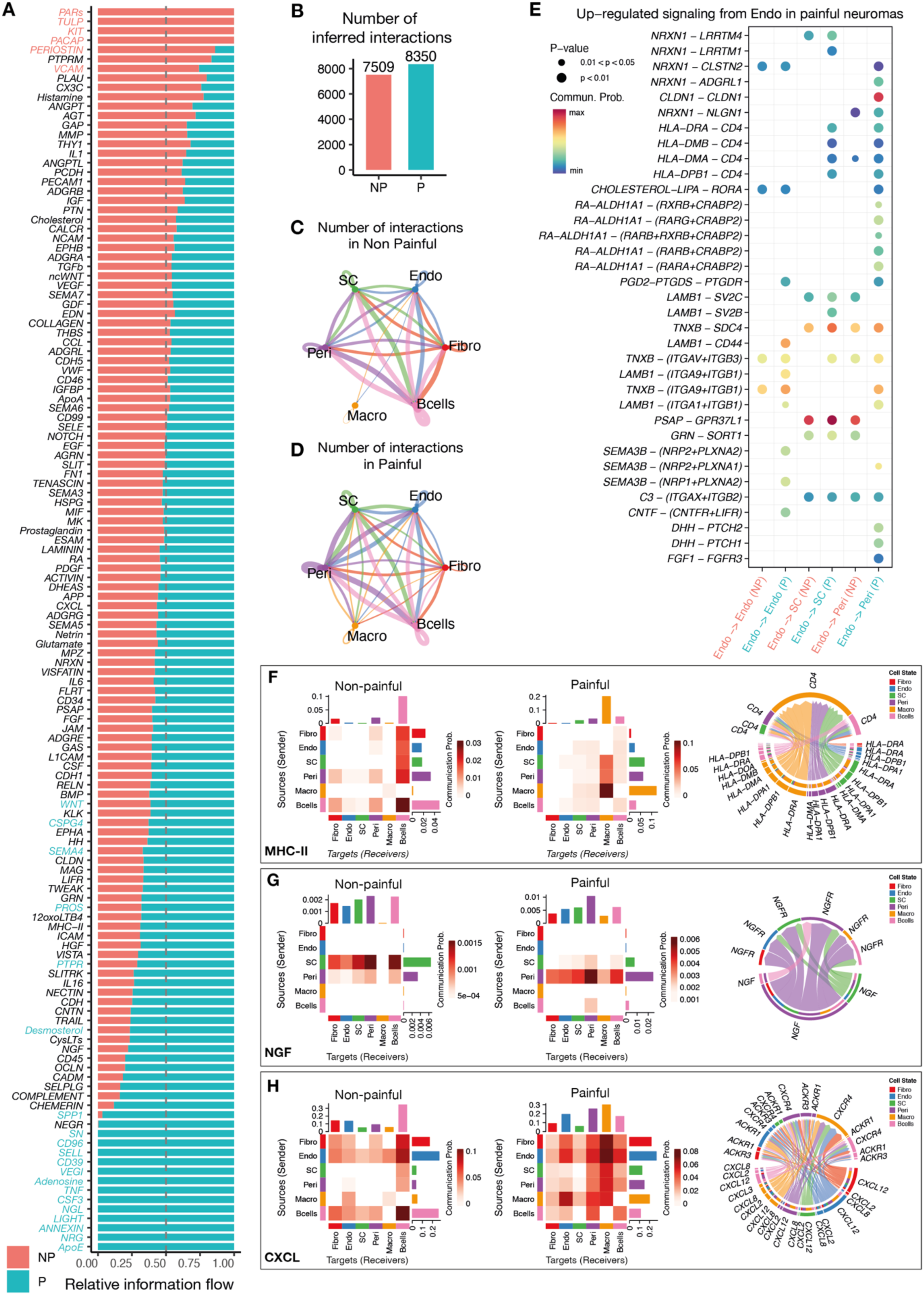
Inferred cell-cell interactions in painful and non-painful neuromas with CellChat v2. A: Barplot displaying the relative information flow of pathways with significant inferred interactions in painful and non-painful neuromas. B: Barplot displaying the number of interactions inferred in painful and non-painful neuromas. C,D: Circle plots displaying the interactions between cell types in non-painful (C) and painful (D) neuromas, where the thickness of the lines is proportional to the number of interactions with a cell type. E: Bubble plot displaying interactions between ligand-receptor pairs that are differentially expressed in painful and non-painful neuromas involving endothelial cells, where the size and colour of each circle represents the p-value and the likelihood of the interaction, respectively. F, G, H: Heatmaps showing the communication between cell-types in painful and non-painful neuromas in the MHC-II (F), NGF (G) and CXCL (H) pathways, where a darker shade of red represents a higher communication probability, as displayed in the scale (note that the scale is different for each heatmap). The chord plots represent the ligand-receptor pairs exhibiting significant interactions in the painful neuromas.

By examining individual pathways, changes in the contribution of cell types can be identified: MHC-II signalling is increased in painful samples (Figure 5.F) where MHC class II ligands are more likely to interact with CD4 expressed within the macrophage cluster rather than in the B cells cluster (Supplementary Fig 2.A); NGF signalling is also increased in painful samples where the main source is the perineural cluster, rather than Schwann cells (as for the non-painful neuromas), (Figure 5.G, Supplementary Fig 2.B) and CXCL signalling displays an increased expression of ligands and receptors in perineurial, macrophage and endothelial clusters (Figure 5.H, Supplementary Fig 2.C).

### HLA-A protein is expressed in human neuromas and expression in nerve fascicles correlates with symptoms of pain

Due to the marked over-expression of *HLA-*A in painful versus non-painful nerve fascicles, detected with spatial transcriptomics, HLA-A protein expression was further investigated by immunohistochemistry. HLA-A strongly co-localised with UEA-I (Vector Laboratories, FL-1061), a leptin that binds to glycoproteins expressed on the surface of endothelial cells (Figure 6.A). HLA-A positive staining was closely associated with nerve fascicles marked by TUJ1 (Figure 6.B), as well as CD45^+^ immune cells, with some cells displaying co-localization (Figure 6.C). The expression of HLA-A was investigated in a total of 9 samples, as indicated in Table 2, including 4 non-painful and 5 painful ones. Patients in the non-painful group reported an average pain VAS score of 1.75, while patients in the painful group had VAS scores ranging between 43 and 100, with an average of 68.2. There was a significant difference in the pain VAS score between the high and low pain groups (p < 0.001). The area of positive HLA-A signal in areas associated with TUJ1 staining was significantly correlated with symptoms of pain (Pearson r=0.69, p<0.05, Figure 6.D-E).

**Figure 6.**
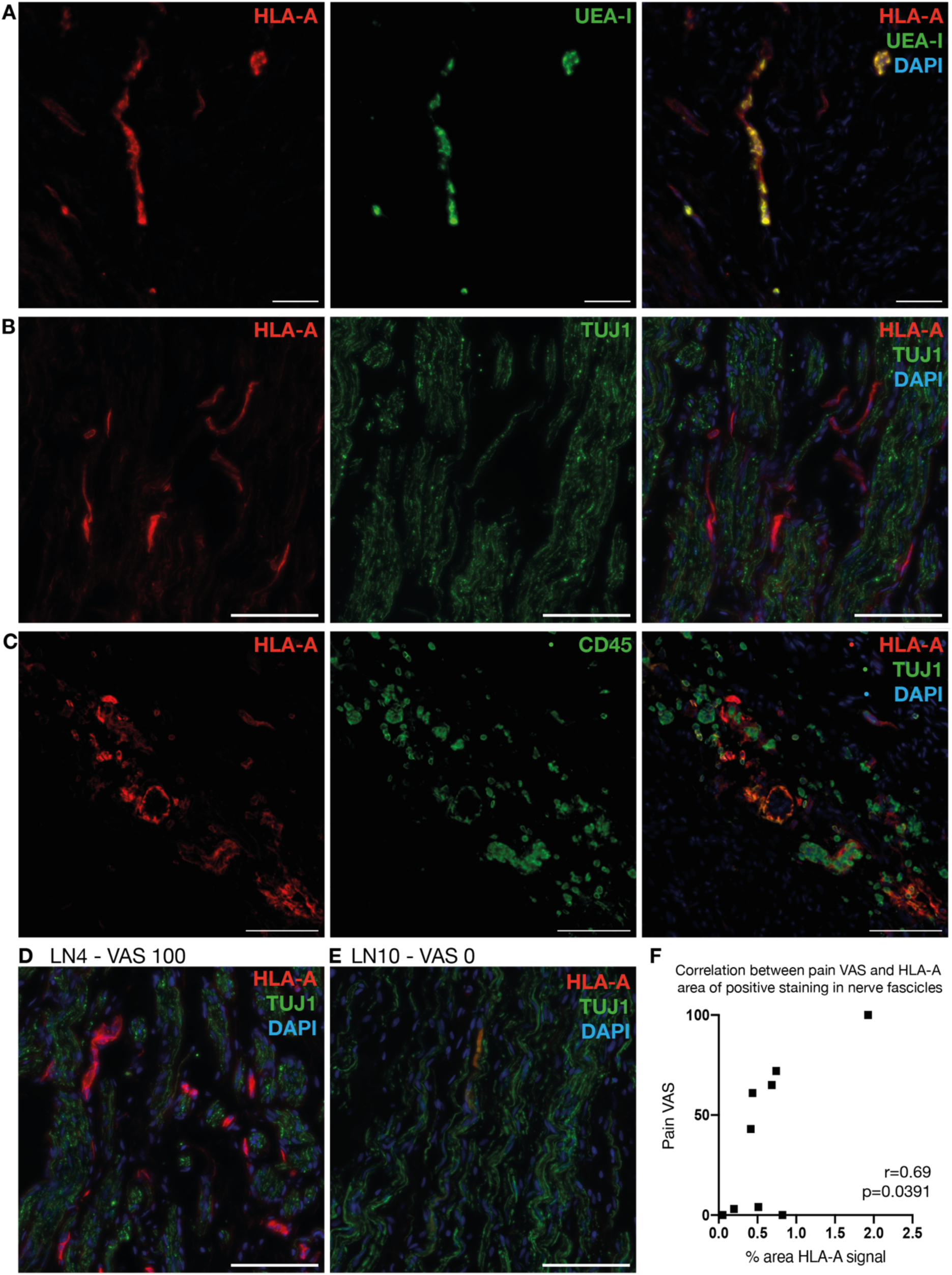
HLA-A immunohistochemistry in human neuromas. A, B, C: Representative sections of HLA-A (Cy3) dual-labelling with the endothelial cell marker UEA-I (FITC, A), the axonal marker TUJ1 (FITC, B), and the immune cell marker CD45 (FITC, C) in samples of human neuromas. Scale bar: 100 μm. D, E: Representative sections of HLA-A (Cy3) signal within nerve fascicles, marked by TUJ1 labelling (FITC) in a painful (LN4, D) and non-painful (LN10, E) sample of human neuroma. Scale bar: 100 μm. F: Correlation between the area of HLA-A positive staining associated with nerve fascicles (identified with TUJ1 labelling) and the intensity of pain reported by the patient. Pearson r= 0.69, p=0.0391.

## Discussion

In order to gain new insight into the molecular landscape underlying neuropathic pain, the cellular composition and the transcriptional landscape in human trigeminal nerves in health and injury were comprehensively characterised using single nuclei RNA sequencing, spatial transcriptomics, RNAscope and immunohistochemistry. The comparison of the transcriptional landscape of human neuromas with and without pain led to the identification of several differentially expressed genes that might play a role in pain development and maintenance.

Among the most highly differentially expressed genes in painful and non-painful neuromas, several are involved in the immune and inflammatory response. *HLA-A*, a member of the major histocompatibility complex (MHC) class I antigens, and *NLRC5*, its trans-regulator^63^ were highly upregulated in painful neuromas, and genetic variants in the HLA region have previously been associated with conditions including post-herpetic neuralgia and traumatic-injury induced neuropathic pain^64–66^. *CXCL2* and *CXCL8* were also upregulated in painful neuromas, confirming the growing body of literature that attributes chemokines a role in pain conditions affecting the trigeminal system^67^: in particular, CXCL2 expression is increased in the trigeminal ganglia of rats following infraorbital nerve constriction^68^, while CXCL8 is increased in the serum of patients with migraine^69^ and trigeminal neuralgia^70^. Other genes differentially expressed between painful and non-painful samples include the basement membrane protein *NID2* expressed in non-myelinating Schwann cells, the matrix metalloprotease *MMP19* expressed in profibrotic fibroblasts and the riboflavin transporter *SLC25A3*, which warrant further investigation.

We observed an expansion in the endothelial cell cluster in injured nerves compared to healthy trigeminal nerve roots, displaying an activated phenotype characterised by the expression of inflammatory markers such as *IL-6, SELE* and *ICAM-1*. In painful neuromas, the endothelial cell cluster was more abundant compared to non-painful ones, displaying increased ligand-receptor interactions with Schwann cells and perineurial cells, involving MHC class II molecules, tight junctions and FGF signalling. Additionally, several of the differentially expressed genes identified by pseudo-bulk analysis are expressed in endothelial cells, including *NLRC5, JMJD1C, CXCL2, SLC25A3, ADGRG1* and *TNFAIP3*. Endothelial cells might play a role in establishing a pro-inflammatory environment by enabling the influx of pro-inflammatory mediators and immune cells, which might contribute to pain chronification following nerve injury.

Previous work on human lingual neuromas investigated correlations between the expression of RNAs and proteins and symptoms of pain, focusing on ion channels including Na_v_1.7, 8 and 9, TRPA1, TRPV1 and the P2X7 receptor^9,71–76^. Among the ion channels investigated, only Na_v_1.8 exhibited a correlation with symptoms of pain^9^. On the other hand, the investigation of miRNA expression in neuromas identified several candidates which are dysregulated in painful neuromas and putatively target interleukin and chemokine receptors^75^.

Other studies on human tissue have also identified inflammatory mediators to be linked with the presence of neuropathic pain in human samples. Ray, et al. ^77^ investigated the transcriptome of 50 human dorsal root ganglia derived from thoracic vertebrectomy, where a portion of the samples was associated with neuropathic pain. Despite detecting increased spontaneous activity in the DRG neurons associated with pain, the transcriptional changes detected were mainly linked with inflammation but not with ion channels expression or regulation^77,78^. The most highly differentially expressed genes identified included Oncostatin M and *CXCL2* in males and genes linked with interferon signalling in females^77^. In human Morton’s neuromas, the presence of transcriptional signatures characteristic of macrophages and B cells was higher than in control samples and immunohistochemical analysis confirmed the increased density of CD163^+^ macrophages in neuromas compared to healthy nerve, while the density of CD68^+^ macrophages correlated with burning pain symptoms^10^.

The investigation of pathologically relevant human tissue is highlighting the prominent role that inflammation and the immune system play in neuropathic pain. Evidence from gene association studies identified the importance of several loci in the HLA region that contribute to neuropathic pain susceptibility or confer protection following peripheral nerve injury^79^, complex regional pain syndrome^80^, diabetes^81^ as well as in rat strains with different susceptibility to the development of mechanical allodynia following nerve injury^82^. The functional effects of the variants identified haven’t been investigated, however one might hypothesise that variation in the HLA region might be indicative of hyperactivity in the immune system where tissue injury triggers exaggerated immune responses that lead to maladaptive changes in the nervous system. This might occur via upregulation and increased antigen presentation, heightening immune activation, as well as by the potential direct activation of nerve fibres by HLA antigens.

Overall, a picture is emerging where the immune system plays a significant role in whether an individual is likely to develop neuropathic pain following nerve injury. Further work to confirm the findings in this study on a larger number of samples is required, as well as functional investigation of the differentially expressed genes and whether they play a causative role in neuropathic pain development and maintenance and might act as targets for novel therapeutics, or if they can be detected systemically and employed as biomarkers for patient stratification.

In summary, the work presented here provides a unique atlas describing the cell-type composition and transcriptional signatures in human healthy and injured trigeminal nerves, with and without neuropathic pain. The atlas highlights the role of endothelial cells in neuropathic pain, which display a pro-inflammatory phenotype and are more abundant in neuromas, particularly in the samples linked with pain, compared to healthy nerves. Additionally, differential expression analysis indicated over-expression of *HLA-*A and *NLRC5*, involved in antigen presentation, and the chemokines *CXCL2* and *CXCL8*. Protein expression of HLA-A was localised to blood vessels and exhibited a correlation with symptoms of pain. These findings contribute to the idea that inflammation is a prominent aspect of neuropathic pain, with increased antigen presentation and chemokine signalling in painful neuromas. Further work is needed to identify whether genetic factors might predispose the immune system to a heightened response to injury and result in higher inflammation and pain chronification. Nevertheless, the atlas will represent a precious resource for the pain community in order to advance the understanding of neuropathic pain mechanisms, translate findings from animal models and develop novel therapeutics.

## Methods

### Human Lingual Neuromas

Lingual neuromas were obtained from patients attending the trigeminal nerve injury clinic at the Charles Clifford Dental Hospital, Sheffield, UK. Neuromas were collected during nerve repair surgery, carried out by Dr. Simon Atkins. All neuromas were collected with the informed consent from the patients, in accordance with ethical approvals received by the NHS Health Research Authority (HRA) and Sheffield Teaching Hospitals (STH) (19/SC/0308 STH20664). Clinical information including patients’ age, sex and pain history was recorded preoperatively and anonymised.

After surgical removal, the samples were cut longitudinally: one half, intended for single nuclei RNA sequencing, was immediately flash frozen in liquid nitrogen and stored at -80°C, while the other half was fixed for spatial transcriptomics, immunohistochemistry and RNAscope. Fixation was performed overnight with Zamboni’s fixative (0.1 mol/L phosphate buffer, pH 7.4, containing 4% paraformaldehyde and 0.2% picric acid) at 4°C, then the samples were transferred to 30% sucrose overnight at 4°C, placed in OCT for rapid freezing on a cryostat freezing plate and finally stored at -80°C. The tissue was sectioned in a cryostat at a thickness of 10 μm and placed on Superfrost Plus slides (Epredia, 10149870). Sections were dried for 45 minutes and stored at -80°C.

Based on the pain VAS scores, the neuromas chosen for this study were divided in two groups: ‘painful’ (pain VAS score higher than 40) and ‘non-painful’ (pain VAS score lower than 15). Differences in VAS scores between painful and non-painful groups were evaluated using an unpaired t-test assuming equal variance.

### RNA extraction

Quality control of the samples used for spatial transcriptomics was performed with the RNeasy FFPE kit (Qiagen, 73504). The incubation with the deparaffinization solution was omitted since the samples weren’t embedded in paraffin. The quality of the RNA was assessed using the Bioanalyzer RNA 6000 Pico chip (Agilent, 5067-1513). Samples with a DV200 higher than 50, indicating that more than 50% of the ribonucleic acids are 200 nucleotides or longer, were deemed to be of satisfactory quality to proceed with the Visium protocol.

### Human Trigeminal Nerve Samples

Trigeminal nerve samples were obtained from The Netherlands Brain Bank, Netherlands Institute for Neuroscience, Amsterdam (NBB). All material has been collected from donors for or from whom a written informed consent for a brain autopsy and the use of the material and clinical information for research purposes had been obtained by the NBB.

### Nuclei isolation for snRNA-seq

On the day of isolation, utensils including forceps, scissors and Dounce homogenizers were sterilised and pre-chilled. Surfaces were cleaned with 70% ethanol followed by RNase Zap. Nuclei isolation media (0.25 M sucrose, 150 mM KCl, 5 mM MgCl2, 1 M Tris Buffer pH 8.0) was prepared in advance.

Homogenization buffer was made freshly on the day by supplementing the nuclei isolation media with 0.1 mM DTT, cOmplete protease inhibitor, 0.1% Triton-X and 0.2 U/μL RNA inhibitor. Wash buffer was made freshly on the day by supplementing the nuclei isolation media with 1% BSA and 0.2 U/μL RNA inhibitor. The procedure was performed on ice, using low retention pipette tips and soft touch pipettes to prevent nuclei disruption and maximise recovery.

Briefly, the tissue was chopped up into smaller pieces (<1 mm) using sterile scissors in 2 mL of homogenization buffer on ice. Up to four samples were processed in parallel. The homogenate was transferred using a wide bore pipette tip to a glass Dounce homogenizer, further homogenized with a pestle for 15 strokes and left to incubate for up to 2 minutes on ice. The homogenate was filtered through a 70 μm strainer and centrifuged at 800 x g for 7 minutes. The supernatant was discarded, the nuclei were resuspended in wash buffer, centrifuged again, and resuspended in 1 mL 4% formaldehyde fixative solution and fixation buffer (10X Genomics, PN 2000517). A small aliquot of the sample was used to count the nuclei using trypan blue staining and assess their integrity and the presence of debris and clumping.

### snRNA-seq library preparation and sequencing

The samples were prepared for single nuclei RNA-sequencing with the Chromium Single Cell Fixed RNA Profiling for Multiplexed Samples kit (10X Genomics, PN-1000456) according to the manufacturer’s instructions. Following overnight fixation (16-17 h), the nuclei from each sample were hybridized with barcoded probes targeting the whole human transcriptome. Two barcodes were used for each sample, targeting to recover 16,000 nuclei per sample. Following the Gel Beads-in Emulsion (GEM) generation^83^, the left-hand and right-hand probes were ligated and the barcoded primers on the gel bead were hybridized to the probes. The probes were extended to include the unique molecular identifier (UMI), the 10x GEM barcode and a partial TruSeq 1 sequence for Illumina sequencing. Library preparation and sequencing was performed by the Genome Center at the University of Texas at Dallas. Libraries were quality controlled by verifying optimal size using the High Sensitivity DNA Agilent Bioanalyzer kit and sequenced on an Illumina Nextseq 2000.

### snRNA-seq data processing and analysis

The reads were demultiplexed, converted to fastq files, aligned to the human genome reference GRCh38 and counted using the Cellranger software provided by 10X Genomics. The h5 files containing the feature barcode matrices were processed with Cellbender to remove ambient RNA^84^. The parameters used for Cellbender analysis are included in Supplementary Table 4. The pipeline was run on a GPU hosted by the Stanage high performance computing clusters of the University of Sheffield.

Downstream data analysis was performed on R (4.2.3) with Seurat (4.9.9), the parameters used for data filtering and integration are included in Supplementary table 4. ScDblFinder (1.12) was used to remove doublets from this dataset with default parameters, barcodes with a doublet score higher than 0.5 were removed from the dataset^85^. Further filtering was performed removing barcodes with fewer than 500 UMIs, 250 genes, a mitochondrial RNA percentage higher than 5% and a novelty score (the logarithmic ratio of the number of genes and the number of UMIs detected, indicating the complexity of the RNA species detected) less than 0.8.

The data was normalized with Seurat’s SCtransform, with the method “glmGamPoi”. After an initial exploratory analysis of the data, a subset of cell types was identified to be specific to the trigeminal nerve samples, such as a small number of astrocytes, oligodendrocytes and meningeal fibroblasts, or to the neuromas, such as salivary gland cells or myocytes. To retain the cellular heterogeneity of the different sample types, reciprocal PCA (rPCA) was chosen as the integration method, as it results in fewer overlaps between two datasets following integration, enabling the identification of cell types that are unique to each sample type^86^. The rPCA workflow involved selecting 3000 integration features on the SCT transformed objects, which are used as anchors to prepare the SCT integration. PCA analysis is performed on the object split by the sample type, and the integration anchors are identified with the reduction “rpca” using 30 dimensions. Finally, the data is integrated using the previously identified anchors with 30 dimensions, with the normalization method “SCT”.

The standard Seurat workflow was performed for downstream analysis, involving scaling the data, running the UMAP dimensional reduction technique (30 PCs), finding nearest neighbours (30 PCs) and clusters at resolution 0.5. The clusters were annotated using marker genes obtained from the literature, summarised in Table 2.1. Markers for each cluster were calculated with the function FindAllMarkers from Seurat. One cluster was removed as it exhibited high expression of *NEFL, TAC1, CALCA* and other neuronal genes and was deemed to primarily contain ambient RNA genes deriving from multiplexed samples which were run on the same chip.

### Visium Spatial Transcriptomics

Spatial transcriptomics was performed with the Visium Spatial Gene Expression for FFPE kit for the Human Transcriptome (10X Genomics, 1000338). Briefly, fixed-frozen sections of human neuromas embedded in OCT were sectioned at a thickness of 7 μm, placed on the capture areas of Visium slides and dried for 45 minutes. Deparaffinization was omitted and the sections were stained with haematoxylin and eosin (H&E) following manufacturer’s instructions. The slides were mounted with 100% glycerol and cover-slipped without sealing. The tissue sections were imaged through the glass slide with a Leica Thunder DMi8 inverted microscope equipped with a FlexaCam C1 Camera system at 10X magnification. Tile-scans were merged using the Leica Application Suite X (LAS X) software. The coverslip was removed and the slide was placed in the Visium cassette. Decrosslinking for 1 hour at 70°C with TE buffer was performed to reverse the formaldehyde bonds and expose the nucleic acids. The sections were hybridised overnight with probes targeting the whole human transcriptome: 19,144 genes are targeted by the probe set, 17,943 genes without off-target activity are filtered and present in the final output. After probe hybridization, stringent washes with 2X SSC were performed to remove non-specifically bound probes. Then, the left-handed and right-handed probes were ligated with a ligase enzyme and the RNA was digested to allow the release of the probes, which were captured by the oligos present on the surface of the Visium slide. The probes were extended to incorporate the unique molecular identifier, the spatial barcode information and the TruSeq Read 1 index. The extended probes were eluted in 0.8M KOH and transferred to PCR tubes. Quantitative PCR was performed on 1 μl of each library with the KAPA SYBR FAST qPCR Master Mix (Merck, KK4600) to estimate the number of PCR cycles required to amplify each library. Library amplification was performed with a unique 10X Sample Index from the Plate TS Set A (10X Genomics, PN-1000251) for each sample. Short fragments were removed with SPRIselect magnetic beads (Beckman Coulter, B23317). Quality control was performed with the High Sensitivity DNA kit (Agilent, 5067-4626) on a Bioanalyzer. The libraries were sequenced on a NovaSeq 6000 S4 platform with 2×150 cycles (PE150) by Novogene Co., Ltd.

### Visium data processing and analysis

Sequencing files in fastq format were processed with Spaceranger (2.1, 10X Genomics) to align the reads to the human reference assembly GRCh38 and process H&E images to align the fiducial frame and detect the tissue. Downstream processing was performed with the R package Giotto (3.3.1): data from each section was merged, genes were filtered so that the expression threshold would be 1 and they were detected in a minimum of two spots, while spots with less than 100 genes were excluded. Each sample was normalised with a scale factor of 6000. PCA was calculated using the highly variable features expressed in more than 3% of spots with a mininum average detection threshold of 0.4. The sections were integrated using the Harmony method, UMAP and nearest network were calculated to perform Leiden clustering at resolution 0.4 with 1000 iterations. Marker genes for each cluster were calculated with the scran method on the normalized gene expression values.

### Differential abundance analysis

Differential cluster abundance was calculated with edgeR (4.2) using the quasi-likelihood negative binomial generalized log-linear model^60,61^. An EdgeR object was created with the function DGEList, the design was formulated to include the pain status with a blocking factor as the sample of origin of each replicate. The dispersion was estimated for each cluster with estimateDisp. Differences in abundance were tested with glmQLFTest. The logFC, nominal P value and BH-adjusted P value for each cluster are reported in Supplementary table 3.

### GO analysis

GO analysis was performed with topGO (2.56) using the significant marker genes (p<0.001) for each cluster. All genes in the spatial transcriptomics dataset were used as background. The GO database was created with biomaRt (2.60), using the “hsapiens_gene_ensembl” as database. Genes were annotated for their biological process and associated gene ontology terms. GO analysis was run with the topGO function runTest, selecting the “classic” algorithm with “fisher” statistics. P-values were adjusted with the Benjamini-Hochberg method. Enrichment is defined as the number of annotated genes observed in the input list divided by the number of annotated genes expected from the background list.

### PAGE enrichment

Cell-type parametric analysis of gene set enrichment (PAGE) was performed using the marker genes from cell types identified with snRNA-seq within the Giotto suite. PAGE calculates an enrichment score based on the fold change of cell type marker genes for each spot. Markers for each annotated cluster in the snRNA-seq dataset were identified with the scran method on normalised expression values with Giotto. PAGE enrichment was calculated with the Giotto function runPAGEEnrich, using a signature matrix that includes the cell types and the top 10 marker genes for each cell type.

### Differential expression analysis

The feature count matrices in h5 format were loaded in R with Seurat (4.9.9)^87^. The barcodes overlaying nerve fascicles were manually selected on Loupe browser (6.5, 10X genomics) and exported in Seurat. When multiple sections were placed on the same capture area, the nerve fascicles from each section were classified as separate technical replicates. The counts from each section were aggregated for pseudo-bulk differential expression analysis, performed with DEseq2 (1.44)^88^. After performing variance-stabilizing transformation with the vst function from DEseq2, principal component analysis (PCA) analysis was conducted with the package PCAtools (2.16). DE analysis was performed with DEseq2. Nominal P values were corrected for multiple testing using the Benjamini-Hochberg method and the logarithmic fold changes (LFC) were shrunk with the approximate posterior estimation for generalized linear model (apeglm) method to reduce the variance of LFCs caused by low or variable gene counts^89^. Genes with an adjusted p-value lower than 0.05 were deemed to be differentially expressed. The volcano plot and graphs of normalized counts were generated with the R package ggplot. The expression of the top DE genes in the snRNA-seq dataset was visualised with Seurat and ggplot.

### Inference of cell-cell communication

CellChat v2 (2.1.2) was used to investigate changes in cell-cell interactions between the clusters in the Visium data in painful and non-painful samples. CellChat is based on a manually curated database, CellChatDB, which takes into account ligand-receptor (L-R) interactions as well as the presence of co-factors and multimeric complexes. Intercellular communication is calculated based on a mass action model, along with differential expression analysis and statistical tests on cell groups^90^.

The count matrices of all samples, separated by painful and non-painful, and the cluster annotation from Harmony integration performed with Giotto were used to create the cellchat objects, using the identities of Schwann cells (SC), endothelial cells (Endo), perineurial cells (Peri), B cells (Bcells) and macrophages (Macro). The spatial information from json files generated by spaceranger were used to account for the distances between barcodes. Standard cellchat workflow was used for the generation of the cell-cell interaction inference network, using the human ligand-receptor database. Dysfunctional signalling was identified with differential expression analysis. The interactions were visualised with the plotting functions provided by CellChat, including chord plots, circos plots and heatmaps.

### RNAscope

RNAscope in situ hybridization multiplex version 2 was performed as instructed by Advanced Cell Diagnostics (ACD) following the fixed-frozen tissue preparation protocol^91^. Antigen retrieval was performed in a steamer at 95°C for 5 minutes with ACD 1X Antigen Retrieval Reagent. Protease treatment was performed with Protease III at room temperature for 1 minute. RNA quality in all tissues was checked by using a positive control probe cocktail (ACD) which contains probes for high, medium and low-expressing mRNAs expressed in all cells (ubiquitin C, Peptidyl-prolyl cis-trans isomerase B and DNA-directed RNA polymerase II subunit RPB1). A negative control probe against the bacterial DapB gene (ACD) was used to reference non-specific/background label. The following probes were used: RNAscope™ 3-plex *Hs-SOX10-C1*, targeting *SOX10*, and RNAscope™ *3-plex Hs-PTGDS-C2*, targeting *PTGDS*.

### Immunohistochemistry

Frozen sections of human neuromas cut at a thickness of 10 μm and placed on Superfrost Plus slides were thawed and washed in PBS-T (0.2% Triton-X in PBS). The following primary antibodies were used: mouse monoclonal anti-HLA-A (Abcam, ab52922, 1:1000), mouse monoclonal anti-NGFR (Abcam, ab3125, 1:200), rabbit polyclonal anti-PI16 (Atlas, HPA043763, 1:200), mouse monoclonal anti-TUJ1 (BioLegend, 801202, 1:100), rabbit polyclonal anti-GLUT-1 (Abcam, Ab15309, 1:400) and mouse monoclonal anti-CD45 (Abcam, Ab8216, 1:400). Labelling of endothelial cells was performed with fluorescein conjugated Ulex Europaeus Agglutinin I (Vector Laboratories, FL-1061, 1:50). The sections were incubated with 20% Normal Donkey Serum in PBS-T for 1 hour at room temperature. Primary antibodies were diluted in 5% NDS in PBS-T and incubated overnight at 4°C. The slides were washed with PBS-T and incubated with a secondary antibody diluted in 1.5% NDS in PBS-T for 90 minutes in the dark. Sections were mounted in Vectashield Antifade Mounting Medium with DAPI (Vector Laboratories, H-1200-10), cover-slipped and sealed.

### Image Acquisition

Images were acquired using a Leica DMi8 inverted microscope fitted with a Leica K5 sCMOS microscope camera system. The microscope was equipped with a LED 3 fluorescent light source in the 390-680 range, and 4 filter cubes for epifluorescence excitation: DAPI, FITC, Cy3 and Cy5. Tile scans and z-stacks were acquired at 20X and 40X magnification and processed on the Leica Application Suite X (LAS X) software and on Fiji (v2.14).

Images of *PTGDS* and *SOX10* RNAscope were acquired on an Olympus FV3000RS Confocal Laser Scanning inverted microscope at the University of Texas at Dallas. The samples were imaged with 405 nm, 488 nm, 561 nm and 640 nm diode laser lines at 40x magnification. Images were acquired on the Fluoview acquisition software and processed on Fiji.

### Image analysis and quantification

Image analysis was performed in Fiji (2.14) by a blinded investigator. Maximal intensity projections were generated from z-stacks of tile-scans of the tissue sections imaged at 20X magnification. A threshold was selected to isolate areas positive for HLA-A labelling and the percentage of the area within the nerve fascicles, identified by TUJ1 labelling, was recorded. Statistical analysis was performed using GraphPad Prism (10.0.2, GraphPad software, San Diego, CA, USA). Pearson correlation test was used to test the correlation between the area of HLA-A positive labelling with the patient’s self-reported pain VAS scores.

## Data and code availability

Processed snRNA-seq and Visium data will be deposited on the Gene Expression Omnibus (GEO) repository. Raw sequencing data will be deposited on the European Genome-Phenome Archive (EGA). Datasets will be released to the public when the manuscript is published in a peer-reviewed journal. Custom R scripts are available at https://github.com/martina-morchio/pain_human_neuromas.

## Acknowledgements

We would like to thank all participants for taking part in this study. We gratefully acknowledge the staff at the School of Clinical Dentistry, Sheffield, UK, who have helped with the collection of the tissue, in particular Dr Matt Worsley and Katy D’Apice. We would like to acknowledge the contribution of Evgeniya Anisimova by her assistance with the validation of primary antibodies. We thank the Genome Center at The University of Texas at Dallas for the services to support the single nuclei RNA sequencing. We also thank Dwayne Thomas and collaborators at Eli Lilly for their support. This work was supported by the Biotechnology and Biological Sciences Research Council through a Training Grant [grant number BB/T508159/1] and Impact Acceleration Award, Eli Lilly and the Battelle Memorial Institute (Dr Jeff Wadsworth – Battelle Knowledge Exchange Scheme Award 2022).

## Author Contributions

FMB, DWL and MM designed the study. SA performed the nerve repair surgeries and collected the neuromas. MM processed the neuromas, performed RNA extractions, Visium experiments, immunohistochemistry and RNAscope. IS and DT-F performed the nuclei isolation and library preparation for snRNAseq. MM analysed sequencing data. NW and MM analysed the immunohistochemistry experiments. FMB, DWL, ES and TJP supervised the study. MM drafted the manuscript. FMB and DWL reviewed and edited the first draft. All authors reviewed and read the paper.

## Competing Interests

The authors report no conflict of interest.

## Notes

### Competing Interest Statement

The authors have declared no competing interest.

