## Supplementary files for "Investigation of cellular and molecular changes linked with neuropathic pain in healthy and injured human trigeminal nerves"

### Supplementary materials

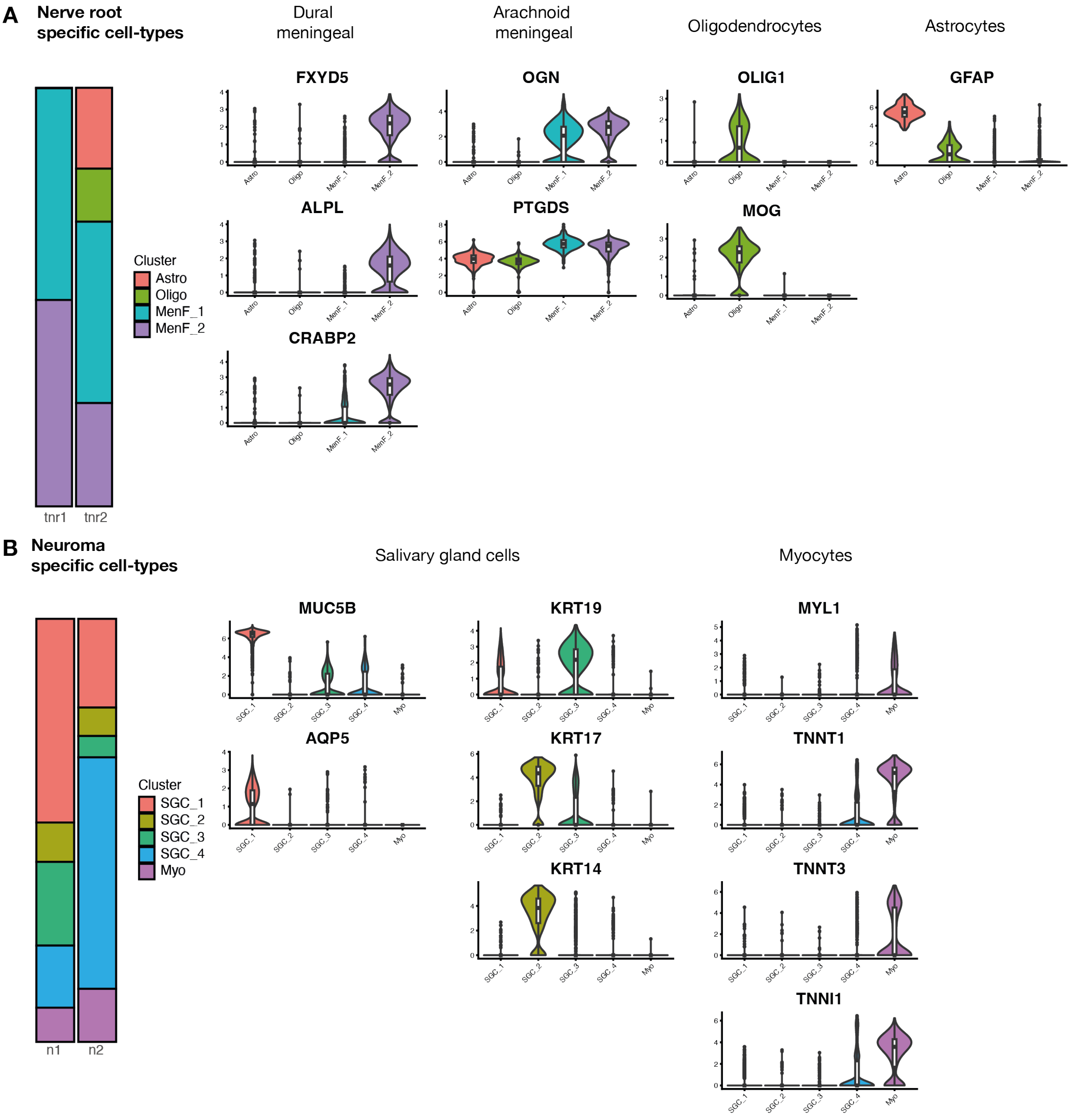


**Supplementary figure 1. Other cell-types identified by snRNA-seq.**

Nerve root (A) and neuroma (B) specific cell types identified by snRNA seq. In nerve root samples, meningeal fibroblasts are identified by arachnoid markers (MenF_1: OGN, PTGDS) and dural markers (MenF_2: FXYD5, ALPL, CRABP2)[1]. Astrocytes are identified by GFAP expression, while oligodendrocytes by OLIG1 and MOG expression[2]. In the neuroma samples, salivary gland cells (SGC) are identified by the MUC5B and AQP5 expression, typically expressed by acinar cells (SGC_1), KRT14 and 17, expressed by basal duct cells (SGC_2) and KRT19 expressed by ductal cells (SGC_3)[3,4]. Myocytes are identified by the expression of troponin genes[5].

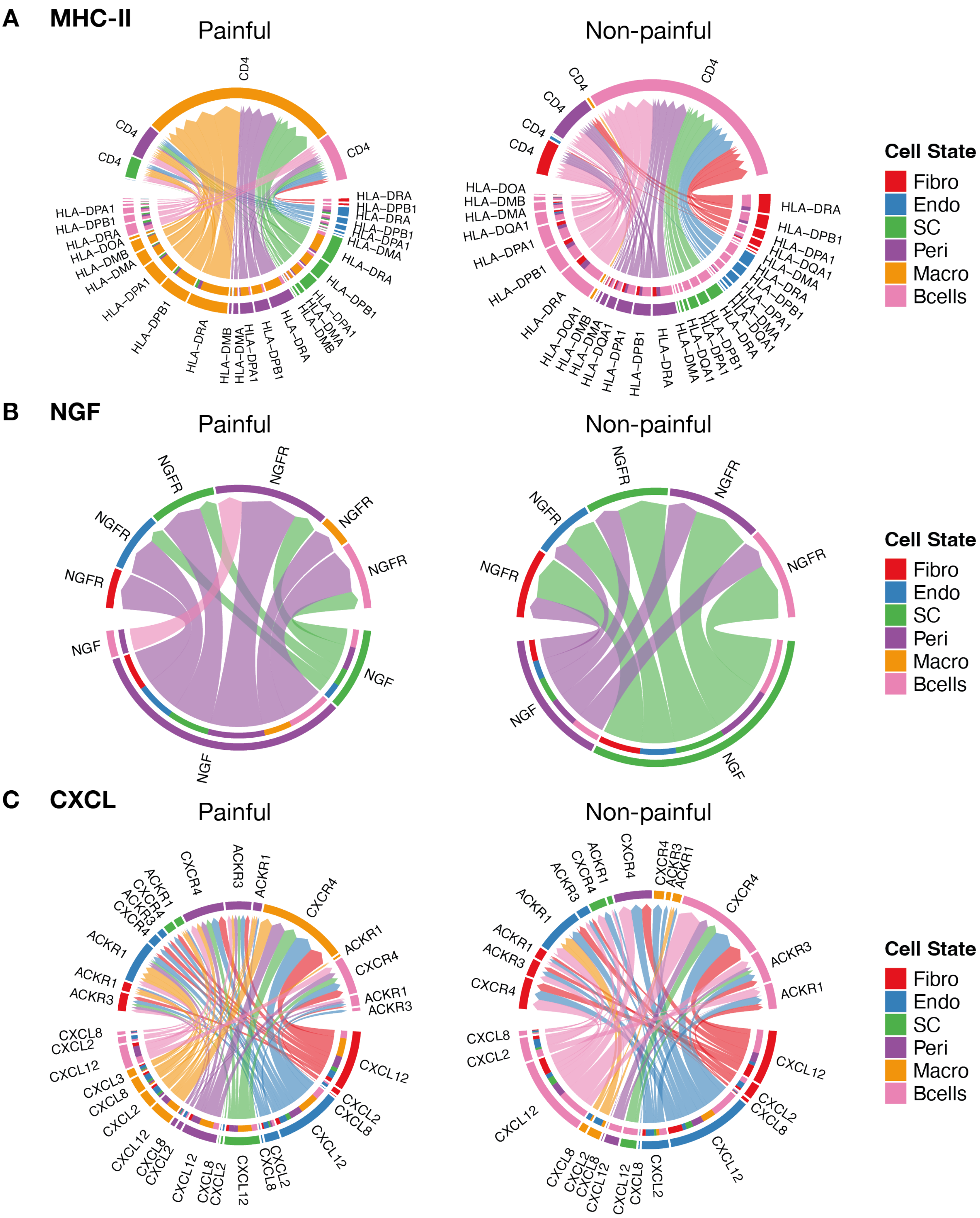


**Supplementary figure 2. Ligand-receptor interactions in painful and non-painful samples in the MHC-II, NGF and CXCL pathways.**

Chord plots displaying the inferred ligand-receptor interactions in painful (left) and non-painful (right) samples for the MHC-II (A), NGF (B) and CXCL (C) pathways. The chords are colour coded by the sender cell-type and directed towards the receiver, whose cell-type is colour coded in the targeted segment. The ligands expressed by the senders are displayed at the bottom, while the receptors expressed by the receivers are displayed at the top.

**Supplementary table 1. Summary of the marker genes used for cell type annotation.**

Markers derived from the literature[1,6-9] used to annotate each cell type are listed.

| Cell Type | Marker Genes |
|---|---|
| General Fibroblasts | DCN, GSN, VIM, COL1A1, FN1 |
| Endoneurial Fibroblasts | OSR2, ABCA8, ABCA9, ABCA10, PLXDC1, COL15A1 |
| Perineurial Fibroblasts | CLDN1, SLC2A1, PTCH1, LMO7, ITGB4, KLF5, NGFR |
| Meningeal Fibroblasts | OGN, PTGDS, FXYD5, ALPL, CRABP2 |
| Endothelial cells | EGFL7, PECAM1, TIE1, EMCN, CDH5, VWF, CLDN5, ECSCR |
| Vascular Smooth Muscle Cells / Pericytes | TPM2, MYH11, ACTA2, MYLK, PDGFRB |
| Schwann cells | SOX10, PLP1, ERBB3, NCAM1, S100B |
| Non-Myelinating Schwann Cells | L1CAM, NRXN1, NCAM1 |
| Myelinating Schwann Cells | MBP, MPZ, EGR2, NCMAP |
| Repair Schwann Cells | NGFR, BDNF, GDNF, ERBB3, SOX2, CADM1, ATF3, RUNX2 |
| Lymphocytes | PTPRC, CD3G, CXCR6, TRAC, CD3E, SKAP1, THEMIS, IL7R |
| Myeloid | AIF1, CD68, MRC1, SIGLEC1, ITGAM, CSF1R |
| Myocytes | MYL1, TNNT1, TNNT3, TNNI1 |
| Oligodendrocytes | OLIG2, OLIG1, MOG, CNP, PLP1 |
| Astrocytes | GFAP |
| Salivary Gland Cells | MUC5B, AQP5, KRT19, KRT7, KRT14 |

**Supplementary table 2. Top 5 marker genes for each cluster and the putative enriched cell-type.**

Each cluster was annotated based on the top differentially expressed genes, where a putative cell type enriched in each barcode was assigned based on gene expression. The number of spots classified as each cell type across all samples is also reported.

| Cluster number | Cluster name | Top 5 DE genes | | | | | N of spots | Putative enriched cell type |
|---|---|---|---|---|---|---|---|---|
| 1 | Fibro | COL1A1 | SFRP2 | FBLN1 | COL1A2 | SFRP4 | 4607 | Fibroblast |
| 2 | Endo | AQP1 | CCL14 | IL6 | TM4SF1 | SELE | 4422 | Endothelial cells |
| 3 | SC1 | HBA2 | MPZ | HBA1 | PMP22 | MBP | 4327 | Schwann cells |
| 4 | Myo1 | MB | TNNT1 | TCAP | CKM | TTN | 4214 | Myocytes |
| 5 | Peri | PTGDS | CLDN1 | SLC2A1 | IGFBP6 | MPZ | 3873 | Perineurial cells |
| 6 | SC2 | MBP | PMP22 | MPZ | PRX | S100B | 3571 | Schwann cells |
| 7 | Myo2 | MB | TNNT1 | TCAP | CKM | TNNI1 | 2936 | Myocytes |
| 8 | SC3 | PMP22 | MBP | MPZ | PRX | APOD | 2882 | Schwann cells |
| 9 | Myo3 | MB | TNNT1 | CKM | TCAP | TTN | 2458 | Myocytes |
| 10 | SC4 | APOD | MPZ | PMP22 | MBP | PRX | 2162 | Schwann cells |
| 11 | Myo4 | ACTC1 | THBS4 | MYLPF | CA3 | COL1A1 | 1937 | Myocytes |
| 12 | SC5 | MPZ | PRX | MBP | PMP22 | S100B | 1549 | Schwann cells |
| 13 | Bcells | IGKC | IGHG2 | APOD | IGHG1 | S100B | 1083 | B cells |
| 14 | SC6 | PMP22 | MBP | MPZ | PRX | S100B | 449 | Schwann cells |
| 15 | Macro | LYZ | MMP9 | SPP1 | LAPTM5 | CHIT1 | 353 | Macrophages |
| 16 | NA | PRX | APOD | MPZ | CNP | MBP | 8 | NA |
| 17 | NA | GPM6B | EGR2 | MPZ | CARD8 | FGFBP2 | 5 | NA |

**Supplementary table 2. Differential abundance analysis**

Differential abundance of cell types between painful and non-painful samples analysed with spatial transcriptomics was calculated with EgdeR using the quasi-likelihood negative binomial generalized log-linear model. For each cell type, the log fold change of relative abundance, the p value and the false discovery rate (FDR) calculated with the Benjamini-Hochberg method are shown.

| | **logFC** | **P-value** | **FDR** |
|---|---|---|---|
| SC6 | 3.26546643 | 3.34E-08 | 5.01E-07 |
| SC2 | 2.51375926 | 7.33E-07 | 4.10E-06 |
| SC3 | 1.14083681 | 8.21E-07 | 4.10E-06 |
| Endo | 0.73955188 | 1.53E-06 | 5.74E-06 |
| Myo1 | -2.7484297 | 2.86E-06 | 8.59E-06 |
| Myo4 | -2.256542 | 0.0002492 | 0.0005429 |
| Myo2 | -2.0639607 | 0.00025335 | 0.0005429 |
| SC4 | 1.04301283 | 0.0005472 | 0.001026 |
| Macro | 1.67939079 | 0.00165775 | 0.00276292 |
| SC5 | -1.07108 | 0.00391552 | 0.00587329 |
| Myo3 | -1.4480828 | 0.00753059 | 0.01026898 |
| Fibro | 0.38803738 | 0.01491649 | 0.01864561 |
| Peri | 0.54213426 | 0.04498707 | 0.05190816 |
| SC1 | -0.2625577 | 0.31190526 | 0.33418421 |
| Bcells | 0.27324782 | 0.58192571 | 0.58192571 |

**Supplementary table 3. Parameters and information for snRNAseq data analysis.**

The table displays information on the snRNAseq dataset, including the metrics calculated by Cellranger in data preprocessing, the parameters used for Cellbender ambient RNA removal, the filtering parameters used in Seurat, as well as parameters used to perform integration and clustering.

| | **N1** | **N2** | **TG1** | **TG2** |
|---|---|---|---|---|
| **Cellranger analysis** | | | | |
| Cells detected | 10,977 | 10,926 | 24,250 | 27,236 |
| Confidently mapped reads in cells | 75.14% | 71.43% | 74.36% | 70.90% |
| Estimated UMIs from genomic DNA | 1.14% | 0.80% | 0.04% | 0.05% |
| Estimated UMIs from genomic DNA per unspliced probe | 6 | 7 | 2 | 2 |
| Median UMI counts per cell | 1,536 | 2,781 | 2,893 | 2,582 |
| Median genes per cell | 983 | 1,550 | 1,773 | 1,666 |
| Median reads per cell | 5,321 | 9,519 | 10,754 | 9,887 |
| Number of reads from cells called from this sample | 79,134,506 | 134,083,140 | 474,861,720 | 426,518,737 |
| Reads confidently mapped to filtered probe set | 94.18% | 95.52% | 90.45% | 87.99% |
| Reads confidently mapped to probe set | 96.01% | 97.21% | 92.44% | 89.74% |
| Reads mapped to probe set | 99.25% | 99.28% | 99.23% | 99.21% |
| Total genes detected | 18,060 | 18,067 | 17,495 | 17,592 |
| **Cellbender parameters and results** | | | | |
| Expected cells | 10000 | 10000 | 20000 | 20000 |
| Total droplets included | 50000 | 50000 | 70000 | 70000 |
| fpr | 0.01 | 0.01 | 0.01 | 0.01 |
| learning-rate | 0.00005 | 0.00005 | 0.00005 | Default |
| epochs | 150 | 150 | 200 | 200 |
| counts in non-empty droplets removed | 3.40% | 3.10% | 5.79% | 8.68% |
| **Filtering parameters in Seurat** | | | | |
| n Cells pre-filtering | 15295.00 | 17392.00 | 25608.00 | 28536.00 |
| nCount_RNA threshold | 500.00 | | | |
| nFeature_RNA threshold | 250.00 | | | |
| log10GenesPerUMI threshold | 0.80 | | | |
| percent.mt threshold | 5.00 | | | |
| nCells post-filtering | 10892.00 | 11193.00 | 16053.00 | 18993.00 |
| n clusters (res= 0.5) | 19.00 | 19.00 | 14.00 | 16.00 |
| **Integration and clustering in Seurat** | | | | |

| | | | | |
|---|---|---|---|---|
| normalization method | "SCT" | | | |
| reduction | "rpca" | | | |
| resolution | 0.50 | | | |
| communities | 27.00 | | | |
| n Cells after cleanup of clusters | 10847 | 11143 | 16047 | 18922 |
